# Mind the alignment gap: a spatial transcriptomics benchmark for scientific coding agents

**DOI:** 10.64898/2026.07.05.736638

**Authors:** Yiqun Chen, Stephanie C. Hicks

## Abstract

Scientific coding agents are difficult to benchmark because many research tasks require executable work yet produce ambiguous or hard-to-verify outputs. Because benchmark construction requires substantial time and resources, automation offers a path to accelerating methods evaluation. We introduce an interactive framework for constructing scientific-agent benchmarks from peer-reviewed papers and diagnosing agent behavior through trace inspection. We apply it as a case study in spatial transcriptomics alignment, constructing 40 tasks from SABench in which agents submit coordinate tables aligning pairs of two-dimensional tissue slices. Across 120 runs and three configurations, we compare a basic prompt, a package-aware prompt, and a full prompt with a prebuilt virtual environment. In this setting, richer package and environment context increased tool exploration but reduced the mean alignment score relative to the basic prompt (0.36 vs. 0.43; 95% CI, [-0.11,-0.03]). Trace inspection showed that added scaffolding often induced unnecessary transformations, fragile package-first workflows, and infrastructure failures. These results illustrate how specialized tooling can alter agent behavior and why scientific-agent benchmarks should evaluate agent traces and the workflows that produce them in addition to the final outputs.

## 1 Introduction

The advance of artificial intelligence (AI) capabilities is rapidly changing scientific practice and data analysis. At the knowledge level, frontier AI systems have excelled on real-world software engineering bug fixes and on graduate-level math and science questions that are difficult even for highly trained humans [1–4]. More importantly for scientific workflows, agentic systems promise a form of “end-to-end” execution: command-line coding agents such as Codex and Claude Code can read and modify codebases, install packages, run experiments, analyze outputs, create visualizations, and produce written summaries from high-level instructions [5–7]. As models and agents continue to grow in capabilities and paradigms, systematic evaluation has become the next bottleneck for further progress.

Early code-generation benchmarks such as HumanEval and MBPP focus on synthesizing short programs that pass unit tests [8, 9], while repository-level benchmarks such as SWE-bench evaluate agents on real GitHub issues with regression test suites [1]. These settings have been central to measuring progress because they admit relatively clear *automated verification*. More recent benchmarks extend evaluation to data analysis, machine-learning (ML) engineering, and scientific workflows: ScienceAgentBench constructs data-driven discovery tasks from peer-reviewed publications [10]; efforts including DSBench, MLAgentBench, MLE-bench, and AutoKaggle target realistic data science and ML pipelines [11–14]; and DiscoveryBench studies whether agents can recover or verify scientific relationships from data [15].

Real scientific research, however, is more iterative and open-ended than most existing benchmarks: tasks are rarely precisely defined, success is multi-dimensional, and ground truth is often ambiguous and expensive to evaluate. Recent efforts have therefore leaned on expert curation, hierarchical rubrics, and bespoke challenge environments [16–18], with curators often prioritizing discrimination and difficulty so that future systems can hill-climb on older ones. We argue this hill-climbing-focused framing should be complemented by a *user-centric workflow* in which scientists inspect what an agent attempted, assess whether the approach was scientifically sound, and iteratively adapt the agent (e.g., via prompt, skill, or harness engineering) to best suit their needs.

This motivates us to consider scientific artifacts such as papers not only as static human-readable documents, but as precursors to active agentic systems. Our work is inspired by many along the same lines: PaperBench evaluates whether agents can replicate conference ML papers by decomposing each replication into many author-informed, gradable subtasks [16]; Paper2Agent takes a complementary direction and converts papers and codebases into interactive agents that expose paper-specific methods as callable tools for language models [19]. We take a focused approach and evaluate the feasibility of converting benchmark papers into sources of executable, scorable tasks.

Our recipe applies broadly to benchmark papers whose input–output structure and evaluation criteria are clearly described; we use papers not as full-paper replication targets, but as blueprints for executable, domain-specific evaluation suites and improvement benchmarks for AI agents. We illustrate the idea using a recent benchmark of spatial transcriptomics alignment methods [20]; this focus on a single analysis task family lets us examine failure modes that simpler multiple-choice benchmarks rarely expose, while leveraging the author team’s background in computational genomics to evaluate method soundness. **Importantly, we do not claim that spatial alignment exhausts scientific agent evaluation; rather, we view it as a concrete case study**, with replication in other domains needed to assess generalizability.

Spatial alignment (SA) of genomics data aims to reconstruct the three-dimensional (3D) molecular architecture of tissues from two-dimensional (2D) slices — such as spatial transcriptomics measurements — using computational techniques. This analysis task is central to spatial biology because integrating multiple tissue slices can reveal 3D tissue organization and molecular interactions that are not directly accessible from individual 2D measurements. Formally, an SA task takes two or more slices with observed spatial coordinates and gene-expression profiles, and asks for a transformation or correspondence map between them. This map should align spatially and molecularly similar regions while accounting for noise, tissue deformation, and biological variation across slices.

Using the spatial alignment cse study, we assess in this paper whether:

1. a computational genomics benchmark paper can be repurposed as an evaluation suite for AI coding agents;
2. current coding agents can leverage specialized computational packages to solve specialized scientific tasks such as aligning spatial transcriptomics data; and
3. explicit prompting and access to specialized packages could lead to better performance.

We find that (1) AI agents produce reasonable and performant baseline heuristics for alignment; but (2) surprisingly, while more carefully crafted prompting steers AI agents away from heuristics toward more specialized packages, this does not consistently improve performance. In particular, even with a virtual coding environment and packages installed, agent performance can degrade due to failed calls, brittle package integration, and internal proxy metrics only weakly aligned with the gold-standard metric.

Given these findings, we advocate that intermediate artifacts — agent traces and logs — should be shared as diagnostic signals for understanding where scientific agents succeed [21], where they fall back on (sometimes surprisingly performant) heuristics, and how future user-in-the-loop scientific benchmarks should be designed.

## 2 Methods

We convert a recent spatial alignment benchmark `SABench`[20] into executable hidden-evaluation tasks for coding agents. Drawing on existing best practices [22–24], we make the public inputs (e.g., genomics data) and an output schema visible to the agents and hide the labels, held-out expression targets, gold coordinates, metric definitions, and scorer metadata from the agents. In our experiments, we evaluate all submissions with the *same* hidden scorer and vary only the support and enviorment available to the agent.

As a proof of concept, we selected 40 of the 295 tasks in `SABench`: we began with a stratified sample across spatial transcriptomics technologies and platforms, omitting tasks whose provided code or task definitions could not be reproducibly reconstructed due to missing intermediate consolidation files or derived assets. For each task, we used a custom script to decompose the task into an agent-facing workspace and evaluation-only files. Specifically, the agent-facing workspace contains the task prompt, paths to the gene expression matrices and coordinates, and the required output schema. The evaluation assets include the “gold” labels (e.g., labeled spatial or anatomical domains across slices), 3D or atlas coordinates, and task-specific scoring criteria. Final scoring is invoked via Python scripts only *after* the agent finishes writing output files.

### 2.1 Agent setup

We consider three execution conditions in increasing order of prompt richness and package availability: Basic, Package-aware (PA), and Full + prior (FPP) (**Table 1**).

**Table 1:** Summary of agent execution conditions.

| Condition | What is provided to the agent |
| --- | --- |
| Basic | Task prompt only; no paper-method context and no prebootstrapped method environments. |
| Package-aware (PA) | Paper-method names and package suggestions in the prompt; the agent installs and probes packages itself, with no prebootstrapped method environments. |
| Full + prior (FPP) | Prebootstrapped method-package environments plus SABench-derived high-level method priors by task family. These priors summarize which paper methods were favored for related task families, but they do not expose hidden labels, held-out expression targets, or scorer outputs. |

Following recent agent benchmarks for computational biology tasks [25], we report runs with Codex CLI with model `gpt-5.3-codex`and reasoning effort set to `xhigh`. The runner uses the Codex approval-bypass mode to avoid repeated human check-ins during autonomous task execution; the exact flag is documented in the appendix.

To assess the variability in agent performance due to LLM stochasticity, we fix the data splits and evaluation schemes and run run three independent replicates for each dataset and execution condition. Each replicate is invoked using the same command in independent workspaces. After the agent completes, a separate evaluation script runs the leakage audit (i.e., whether the ground truth columns hidden to the agent was used in the final submission).

### 2.2 Agent Evaluation

After an agent exits, we run the evaluation script, which is hidden from the agent, on its submitted coordinate files. The evaluation scorer first checks the task contract: each required output file must be present, have the expected schema, and contain finite coordinates. Missing files, schema errors, row-identity mismatches, and non-finite coordinates are treated as failed submissions and assigned a score of zero (out of one).

Valid submissions are then scored with the metric profile specified for that task, following the general rule in SABench [20]: tasks without ground-truth coordinates are evaluated using expression agreement and label or landmark consistency between aligned slices, whereas tasks with hidden gold coordinates additionally report alignment error based on differences between the aligned and true coordinates. We standardize all metrics to lie between zero and one (higher is better) and report the arithmetic mean of all scores as our primary composite score (see **Table S1** for details).

For the subset of tasks where ground-truth coordinates are available, we also report mean absolute error (lower is better) separately, as it ranges from 0 to *∞* rather than being bounded by one. When a task involves aligning multiple slices, we report metrics averaged over consecutive slice pairs. Finally, beyond scorer outputs, we analyze run traces and generated code to characterize agent behavior with LLM assistance. Specifically, we use an independent LLM call to extract failure stages, error messages, package-use signals, and method-use signals from observable evidence.

## 3 Findings

### 3.1 More prompt context increased package use, but not aggregate accuracy

We first checked whether the designed prompt and workspace changes (i.e., Basic, PA, and FPP) actually changed agent behavior. Figure 1 displays the average end-to-end scores (panel A) and observed package usage (panel B) across all tasks. We note that, as expected, giving agents more package-specific context made them more likely to use spatial-alignment packages in their final solutions. FPP, which includes specific package hints as well as prebuilt virtual environments with installed packages, produced the strongest observable package-use signals in their final submissions: PASTE2 in 72% of runs, PASTE in 68%, Spateo in 54%, and SPACEL in 43%. By comparison, PA showed a narrower package footprint, with roughly 48% PASTE use, less than 5% STAligner use, and little meaningful use of other packages. Basic showed almost no package use, except for roughly 20% PASTE calls, likely because PASTE has a simple API and was frequently the first package mentioned in the paper context, and therefore in our prompts.

**Figure 1:**
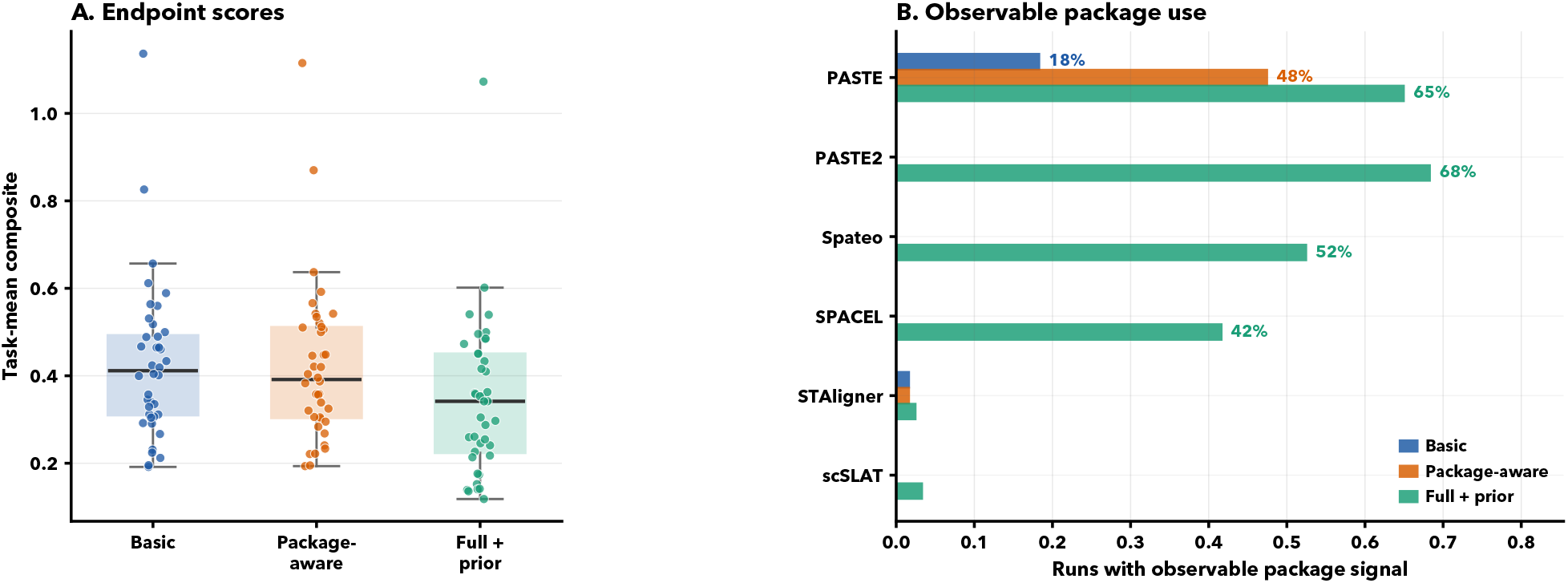
Endpoint and behavior summaries for the 40-task common subset. Left: per-task mean composite scores. Right: observable package-use signals. Full + prior produced the strongest package-use signal, but this behavioral shift did not translate into higher benchmark scores.

The more surprising result is that this increase in package use did not translate into uniformly better performance. In panel A of Figure 1, each point represents one task’s mean composite score across three replicate agent runs. The condition with the *strongest package-use signal*, FPP, had the *lowest average scores*. On the 40-task common subset, Basic achieved the highest mean composite score (0.43, 95% bootstrap CI [0.38, 0.49]), followed closely by PA (0.42 [0.37, 0.48]). FPP was lower (0.36 [0.31, 0.42]). Paired task-level comparisons tell the same story: FPP underperformed Basic by −0.067 (95% paired-bootstrap CI [−0.11, −0.028], Wilcoxon signed-rank *p* = 0.0021) and PA by −0.059 ([−0.11, −0.017], *p* = 0.021), while PA and Basic were not significantly different (−0.0080 [−0.043, +0.029], *p* = 0.48).

We also display the trends of individual tasks (**Figure 2**) to verify that the result is not dominated by a few tasks that happen to perform drastically worse with more package use. We stratified tasks by the Identity baseline, where Identity means submitting the input coordinates unchanged. Easy, Medium, and Hard tasks are defined by tertiles of the Identity composite score, such that a task is considered easier if performing no alignment already yields a relatively high score. We observe that while a few tasks show large variation across conditions, the overall trend declines as more package-specific context is provided. This drop is especially visible on Easy tasks, where some FPP solutions perform no better than a no-op Identity submission.

**Figure 2:**
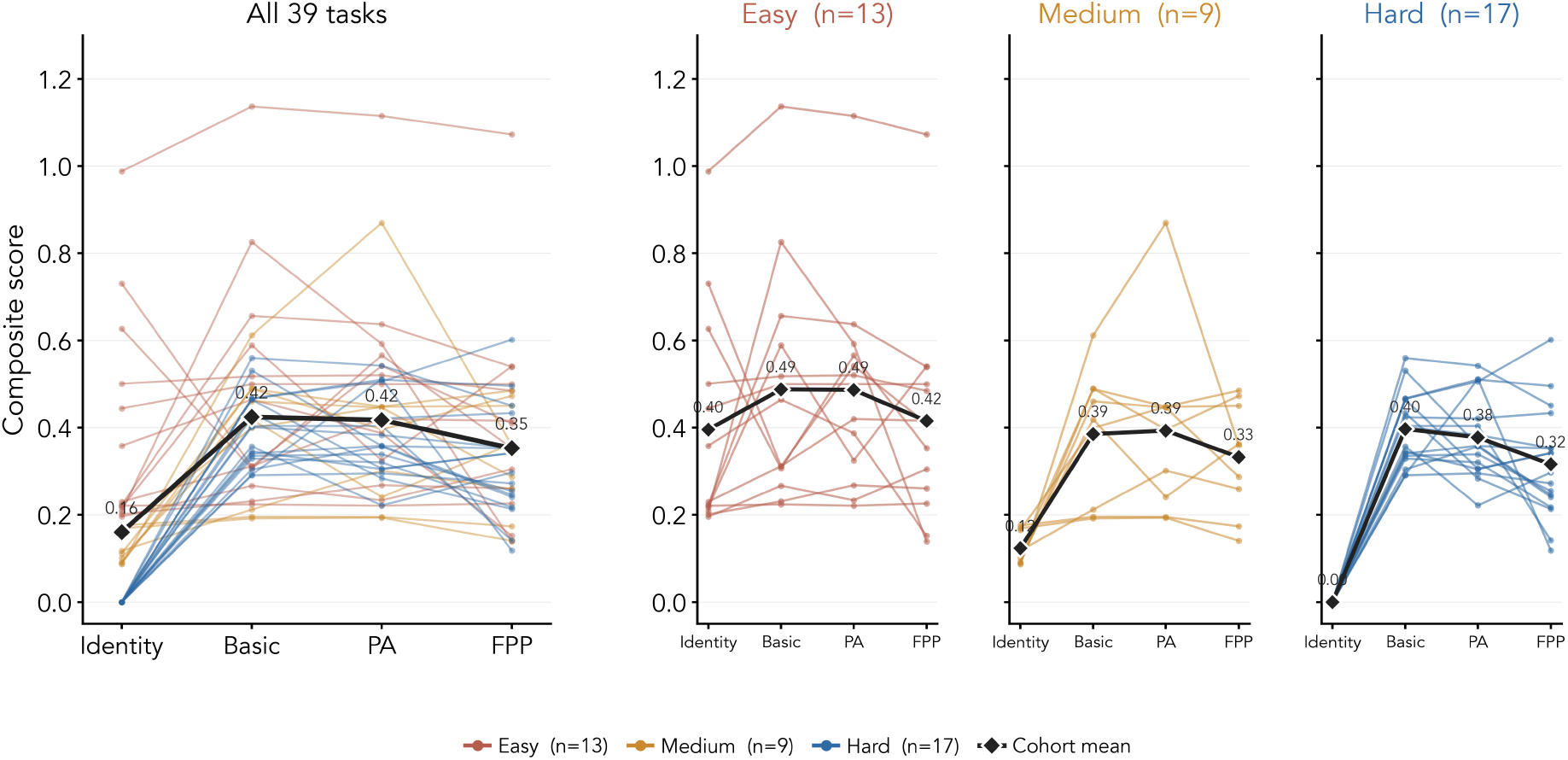
Per-task trajectories on the composite score. Each thin line is one task colored by Identity tier; black diamonds denote cohort means. Left: all 39 scored tasks. Right: the same trajectories faceted by tier. Tertile cutoffs are Hard (Identity ≤ 0.00, *n* = 17), Medium (0 < Identity ≤ 0.18, *n* = 9), and Easy (Identity > 0.18, *n* = 13). PA denotes Package-aware and FPP denotes Full + prior. In every tier, Full + prior ends below Basic.

One possible concern is that the composite score could be driven by only a few metrics that dominate the overall trend. We therefore examined the individual metric families; the full per-metric breakdown is reported in the Appendix. Across all metrics, FPP scores lower than Basic on every individual metric on average. The largest negative differences appear for gene cosine similarity, matching accuracy, and region-grid IoU. This suggests that the aggregate trend is not an artifact of one unusually noisy or unbounded metric.

### 3.2 Why more package use did not lead to higher scores

Since specialized spatial-alignment packages were developed with the promise of better alignment outputs, our finding in Figure 1 is counterintuitive. To understand this apparent contradiction, we inspected the logged search processes of the coding agents as well as their final outputs. The logs reveal that while additional context changed agent behavior, it did not always produce better solutions, for several reasons:

1. **Lighter custom strategies were often competitive**. Under the Basic prompt setting, agents more often wrote custom Python code based on simple geometric ideas, such as Procrustes rotation, to align consecutive slices. On many tasks, such coordinate-only strategies matched or exceeded package-backed solutions, particularly when package performance is sensitive to parameter choices.
2. **Package APIs introduced variance without guarantees**. Invoking a spatial-alignment package requires choosing parameters, handling environment dependencies, and recovering from failures. When these choices were poorly calibrated, the resulting transformations were sometimes worse than simpler geometric alternatives.
3. **Agents could be steered toward worse packages under time pressure**. Different packages performed best on different task families, so providing agents with more package context did not uniformly steer them toward better methods. Under completion pressure, agents sometimes adopted fast heuristics that compromised final solution quality.

This finding aligns with the original benchmarking paper, which proposed that a coarse pre-alignment using manual observation and rigid transformations can substantially reduce computational time and improve accuracy for specialized methods. An extreme case is illustrated by the Identity baseline, which submits the public input coordinates unchanged; a run scoring below Identity has therefore actively degraded an already partially aligned input. On the 39 scored tasks, Basic falls below Identity on 2 tasks, PA on 2 tasks, and FPP on 6 tasks. Two representative cases, SA-004 and SA-014, are illustrated in **Figure 3**. In both, the input coordinates already provide a useful alignment, while agent-selected transformations move slices away from that reference due to poorly calibrated function calls.

**Figure 3:**
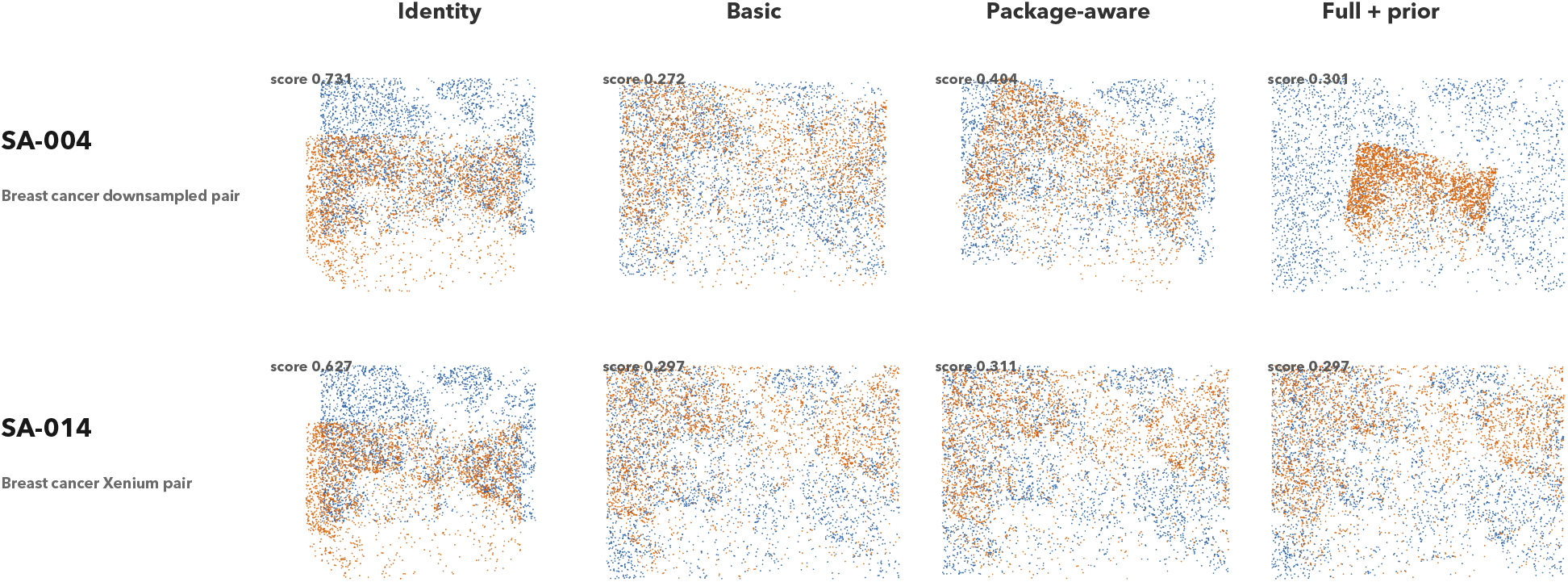
Examples where transforming an already useful input degrades performance. Columns show the Identity reference and representative valid submissions from Basic, Package-aware, and Full + prior. Identity denotes the public input coordinates submitted without transformation; it is a diagnostic reference, not a coordinate-level ground truth. For SA-004 and SA-014, the hidden scorer uses held-out expression and region/landmark agreement rather than a known target coordinate map.

To further investigate failure cases with available ground truth, we inspected MAE for the subset of tasks with hidden gold-standard alignment coordinates. On these five tasks, the advantage of Basic is less clear: differences are often within run-to-run noise, and the lowest mean MAE always comes from either PA or FPP, with FPP best on three of the five tasks. This finding may be confounded by which tasks happen to have gold coordinates (e.g., certain technologies or tissue types may be inherently easier to align). Nevertheless, it flags that results based on gene- and neighborhood-based metrics should be interpreted with some caution, and analyses restricted to tasks with hidden gold coordinates serve as a valuable sanity check. We visualize representative hidden-coordinate tasks in **Figure 4**.

**Figure 4:**
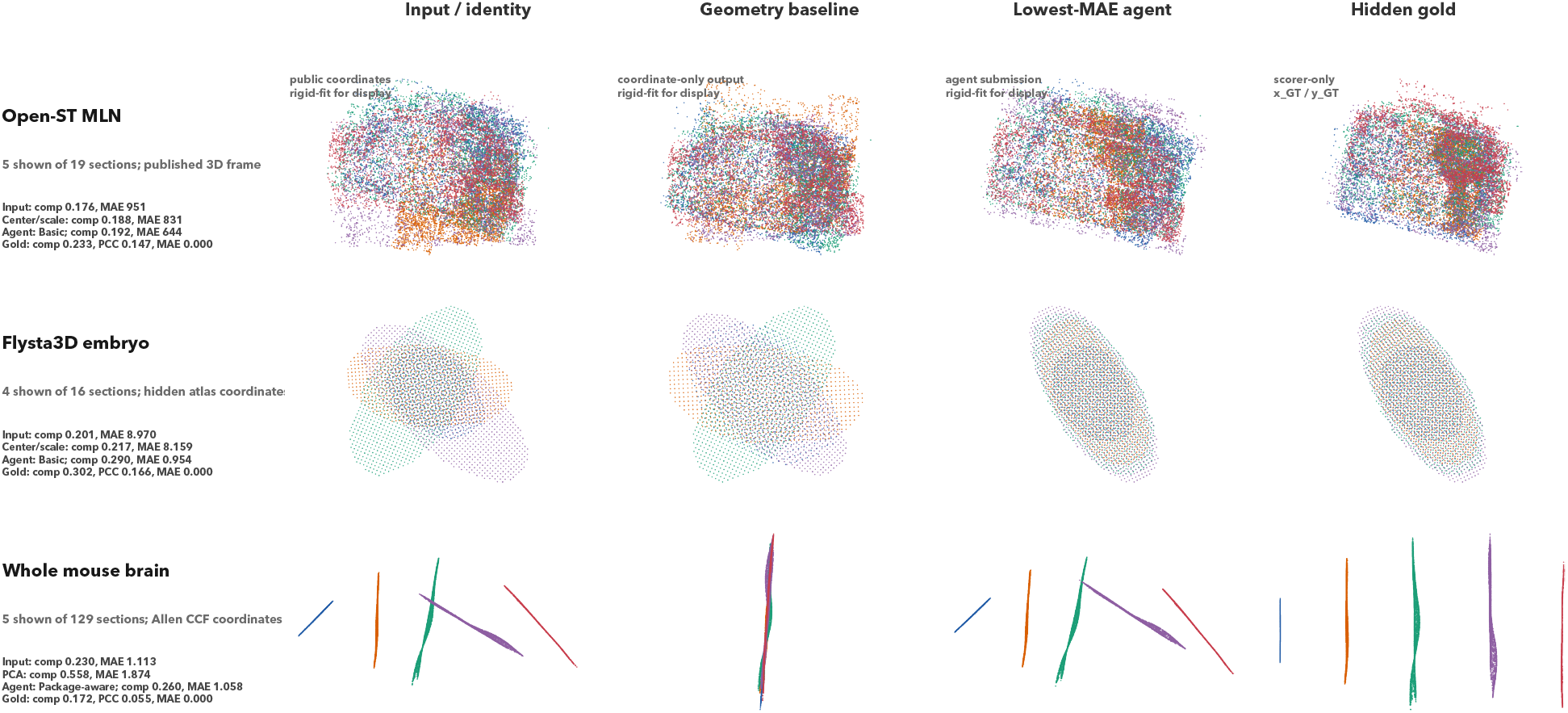
Examples with gold standards. Rows show three tasks with scorer-only 3D or atlas-coordinate references; columns compare the public input, the best deterministic coordinate-only baseline, the valid agent submission with the lowest MAE, and the hidden gold frame. Submitted coordinates are rigid-fitted to the hidden frame for display only. The gold column was never visible to the agent; its MAE is zero by construction, while its displayed composite can differ because the composite excludes MAE and averages higher-is-better expression or label metrics.

Finally, we checked whether the benchmark was dominated by a single package family, which would suggest that a more focused package prompt might resolve the performance gap. We found no such universal winner: rigid or affine custom geometry performs best on some serial-alignment families; Spateo and GPSA are stronger on SCC; PASTE helps in some BCA and AMBC settings; and on DLPFC and MHPR, package and rigid means often cluster closely. The benchmark therefore rewards local method selection over simply increasing package availability (**Table S4**).

### 3.3 Geometric Baselines Inspired by AI Agent Heuristics

The surprisingly strong performance of agent-written geometric solutions motivated us to examine whether Basic agent performance can be *approximated and therefore interpreted* as combinations of simple geometric operations. We synthesized three such methods from agent-submitted solutions. The *center/scale* method translates the moving slice to the reference centroid and rescales it so that the median distance from the centroid matches that of the reference. The *PCA-similarity* method aligns the principal coordinate axes, evaluates the four possible axis sign flips, and keeps the candidate with the largest bounding-box overlap. The *trimmed ICP* method starts from the PCA-similarity and center/scale candidates, iteratively matches nearest-neighbor coordinates by fitting a two-dimensional similarity transform to the closest 85% of matched pairs, and keeps the refinement with the smallest nearest-neighbor distance.

We emphasize that **these probes are not intended as strong scientific methods**; they simply ask whether agents are doing more than basic geometry. On the 38 tasks where they are applicable, center/scale averages a composite score of 0.31, PCA-similarity averages 0.37, and trimmed ICP averages 0.41. On the same tasks, Basic averages 0.42, PA averages 0.41, and FPP averages 0.36. The strongest coordinate-only probe thus nearly matches PA and exceeds FPP.

### 3.4 Illustrative case studies of package wins and losses

To gain visual intuition on when structured package use leads to better performance, we visualize two paired examples in Figure S3. The largest FPP win over Basic is SA-003, a DLPFC slice pair in which one slice’s coordinates were coarsely scrambled during task construction. Basic agents produced inconsistent custom geometry, whereas FPP agents converged on Spateo and PASTE2 alignments. This is the setting where benchmark-derived method context helps: the task calls for the kind of nontrivial transformation that spatial-alignment packages are designed to handle.

By contrast, the largest FPP loss is SA-006, a clean serial alignment of two mouse-brain sections. Basic agents solved the task with short rigid or affine custom solvers. FPP agents instead followed package-backed workflows and submitted alignments that were close in broad structure but misoriented enough to reduce gene-grid and landmark scores. Here, the additional method context pulled agents away from a simple and sufficient solution.

Trace inspection of the logged runs reveals how these outcomes arise. While FPP steered the search process in the intended direction, it also produced longer package loops, more environment failures, and prior-driven submissions—a series of failed package calls would often pressure agents into accepting a potentially suboptimal result rather than recovering with a simpler approach.

**Table 2:** Concrete observable trace vignettes. Short excerpts summarize visible run artifacts; exact longer excerpts are reported in Section E.

| Trace family | Observable excerpt | Process signal |
| --- | --- | --- |
| Baseline insurance | SA-001 Basic: expression-guided transform unstable; one trial hit a runtime failure. | **Valid baseline first**; risky refinement did not break the submission. |
| Slow package fallback | SA-001 Basic: two long-running paste jobs were terminated; validated CSVs remained unchanged. | **Package attempt stalled**; fallback preserved the run. |
| Geometry + expression check | SA-003 Basic: the same transform was best by geometry and expression consistency. | **Geometry was checked against expression** before submission. |
| Large multi-slice execution | SA-020 Basic: all 33 CSVs were written after serial alignment. | **Complete multi-slice output** was validated after a custom solve. |
| Package-backed selection | SA-008 Full + prior: spateo beat a custom rigid fallback by input-only diagnostics. | **Package result was selected by visible diagnostics**. |
| Package error recovery | SA-006 Full + prior: unequal gene sets triggered an intersect/reorder patch. | **Package data-contract issues were patched**, but the re-run remained slow. |
| Provider disconnect after work | SA-105 Full + prior: PASTE2 was still running; a completed spateo submission existed. | **Endpoint failure hid completed work**; provider failure zeroed the run. |

## 4 Discussion and Conclusion

In this work, we constructed a semi-automated benchmark derived from an existing spatial genomics alignment paper and validated its use for evaluating general coding agents on biomedical tasks. Our key finding is that it is much easier to prompt agents to use specialized packages than to use them effectively: more package-oriented prompting increased package probing and code submissions, but agents were often nudged toward poorly tuned variants that scored lower than their own geometric heuristics.

Our study has several limitations that inform future directions. First, all empirical results come from a single model and agent configuration, leaving cross-model sensitivity unaddressed; future work will evaluate behavior across model families and sizes, as well as different coding-agent harnesses (e.g., Biomni, SWE-agent). Second, our findings are restricted to a single benchmark family and a single source paper, so whether the counterintuitive result—that more context can hurt rather than help—generalizes to other domains remains an open question; a natural next step is to extend this framework to additional genomics tasks and source papers, which would also test the generality of the benchmark-construction approach itself. Finally, since agent-written geometric heuristics proved surprisingly competitive, using them as seed or initialization methods, both in spatial transcriptomics and more broadly, is a promising direction for translating our *empirical observations* into *actionable insights* for improving agent performance.

In conclusion, we propose that benchmark papers can be more than static comparisons among human-authored methods and should instead be harnessed as natural evaluation tasks for *AI agents*. Our proof-of-concept spatial genomics application shows that while agents can invoke real packages and sometimes improve task-specific outcomes, added scientific context can also create action bias that lowers aggregate accuracy. Future work should explore whether scientific-agent scaffolds can help agents balance external priors against task-specific evidence, and thereby become more reliable and efficient solvers.

## Appendix

### A Metric families

**Table S1:**
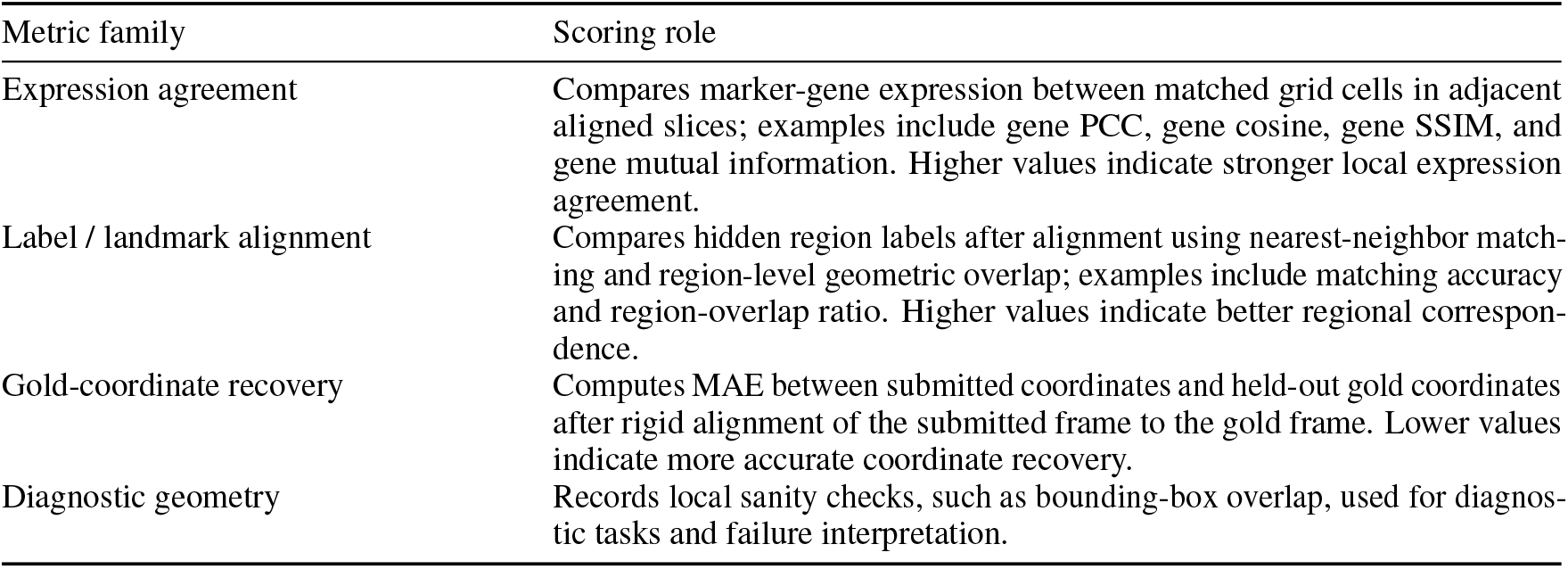
Metric families used by the hidden scorer. The main endpoint is the higher-is-better composite; MAE is reported separately for tasks with hidden gold coordinates. The full task-by-task metric-profile mapping is included in the released task manifests.

### B Task inventory

Table S2 lists all 40 tasks in the common subset with dataset family, scoring profile, slice count, and paper-comparability tier (T1 = exact paper metric table; T2–T3 = same family with caveats; T4 = held-out expression gold only; T5 = local diagnostic).

**Table S2:**
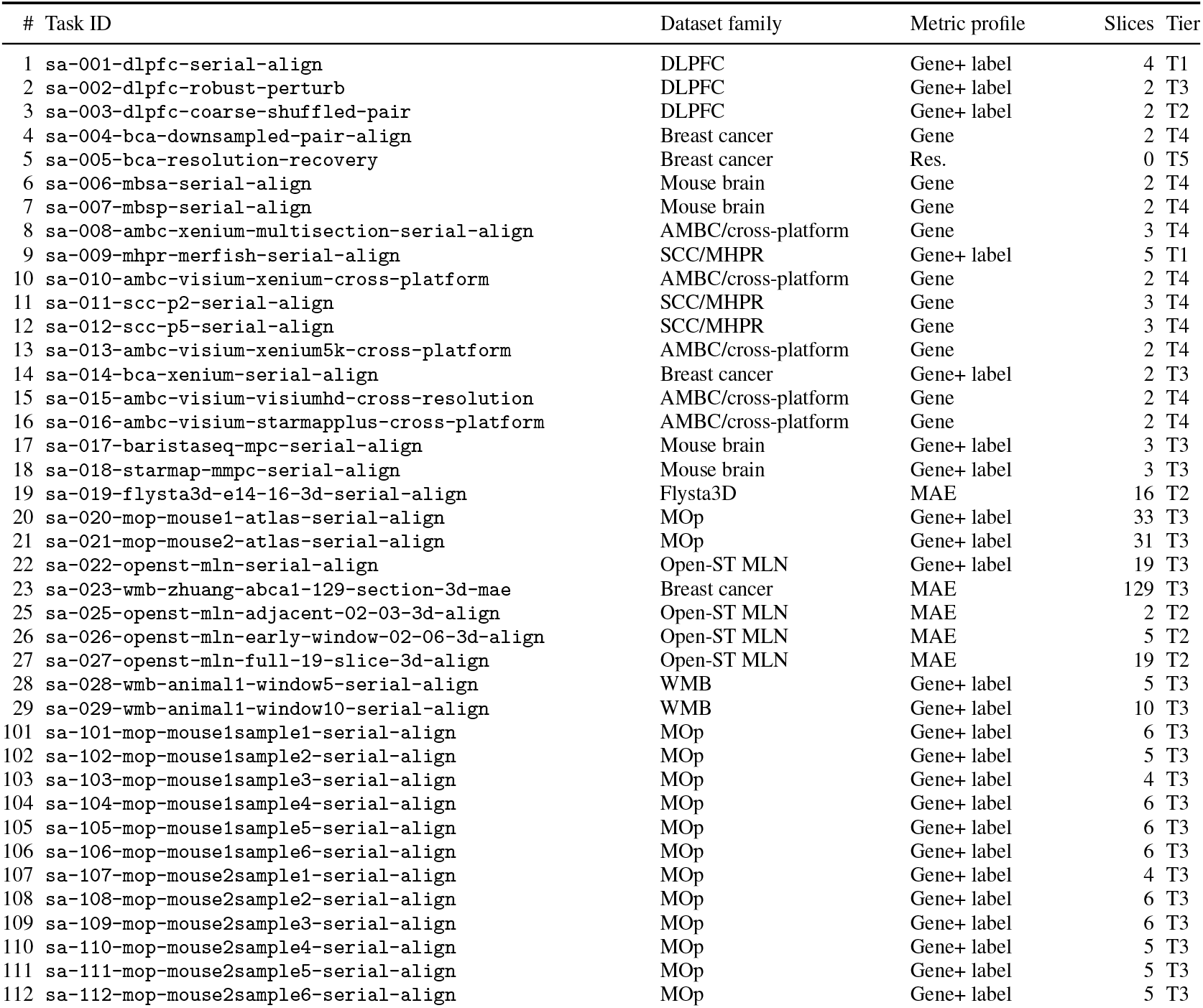
Per-task inventory for the 40-task common subset. List of individual fixed tasks used in benchmark here with corresponding dataset, metric, number of slices, and tiers.

### C Supplementary endpoint results

**Table S3:**
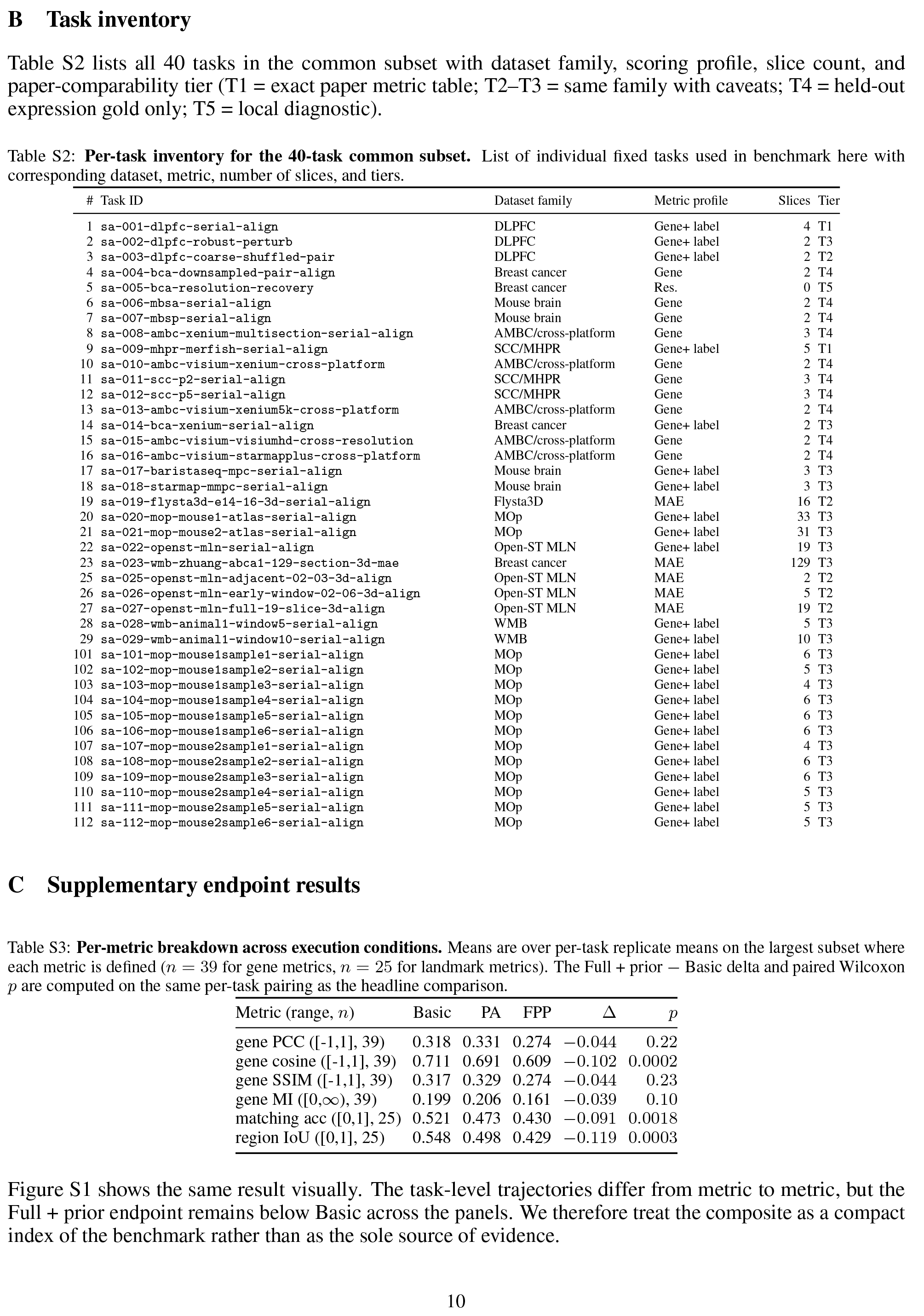
Per-metric breakdown across execution conditions. Means are over per-task replicate means on the largest subset where each metric is defined (*n* = 39 for gene metrics, *n* = 25 for landmark metrics). The Full + prior − Basic delta and paired Wilcoxon *p* are computed on the same per-task pairing as the headline comparison.

Figure S1 shows the same result visually. The task-level trajectories differ from metric to metric, but the Full + prior endpoint remains below Basic across the panels. We therefore treat the composite as a compact index of the benchmark rather than as the sole source of evidence.

**Figure S1:**
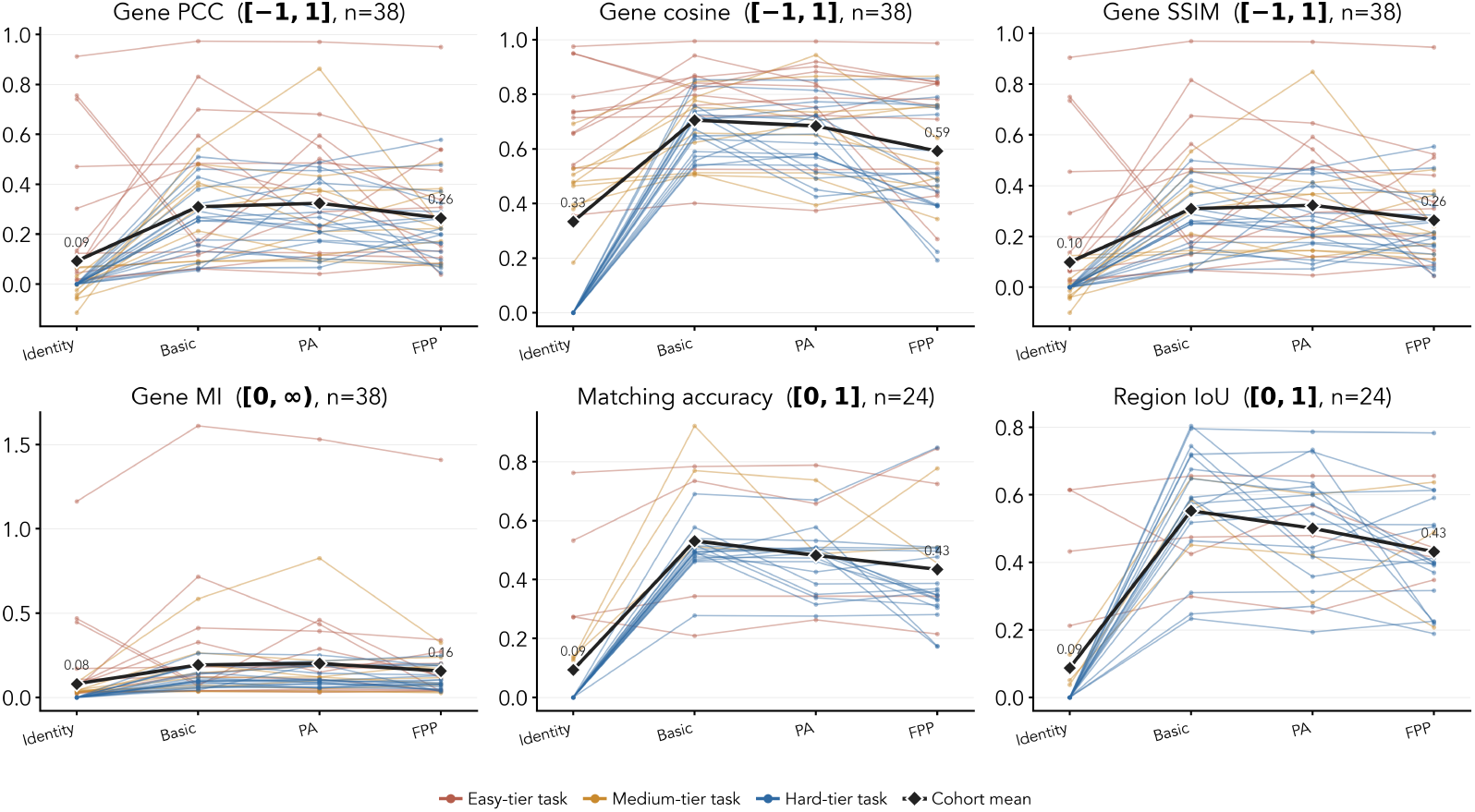
Per-task trajectories by individual metric. Identity tier is assigned once at the task level using composite-Identity tertiles and then carried into each per-metric panel. The Full + prior endpoint sits below Basic on every metric. Axis labels use PA for Package-aware and FPP for Full + prior.

**Figure S2:**
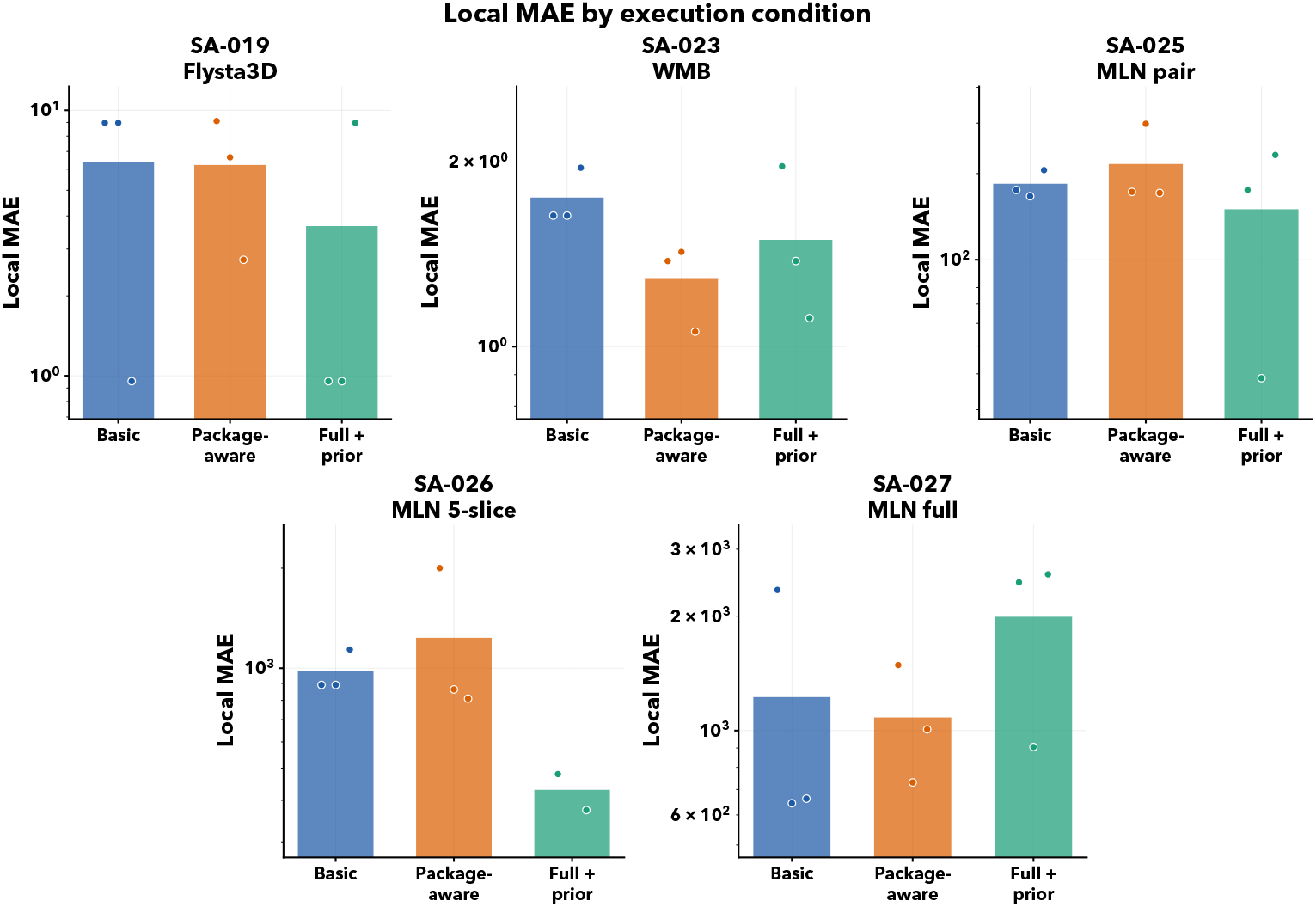
Local MAE comparison across execution conditions for tasks with hidden gold coordinates. Bars show condition means and points show individual runs. Lower is better. MAE is reported separately from the composite because it is lower-is-better and task-scale dependent.

**Figure S3:**
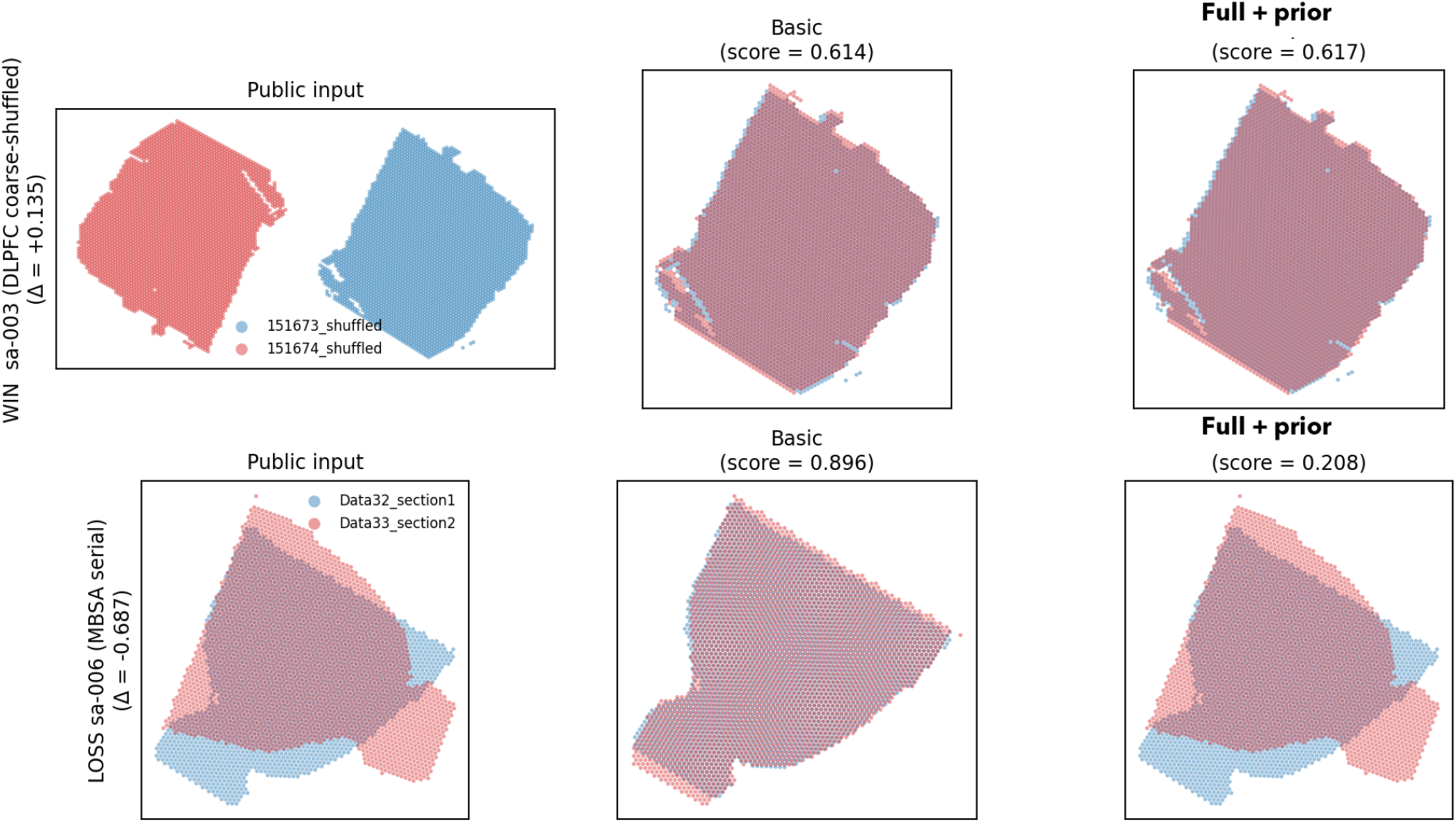
Largest paired Full + prior win and loss against Basic on the 40-task common subset. Each row is one task; columns show the public input coordinates and the highest-scoring Basic and Full + prior submissions. Top: SA-003 is a DLPFC coarse-shuffled pair where spatial-alignment packages help. Bottom: SA-006 is a clean serial-alignment task solved well by short custom geometry but degraded by package-driven alignment.

### D Paper-reference and diagnostic analyses

#### D.1 Additional diagnostics and paper-reference comparisons

The remaining analyses are useful for interpretation but more caveated than the main benchmark comparisons. We therefore treat them as supplementary diagnostics rather than as the primary result.

Paper comparisons require explicit comparability tiers. Only two tasks have exact extracted paper metric tables that align closely with the local task construction: DLPFC serial alignment and MHPR MERFISH serial alignment. Other source-benchmark results provide useful context but differ in metric definition, coordinate scale, or availability of raw per-method tables. Figure S4 overlays agent results with the published SABench method values for the exact direct-table cases.

**Figure S4:**
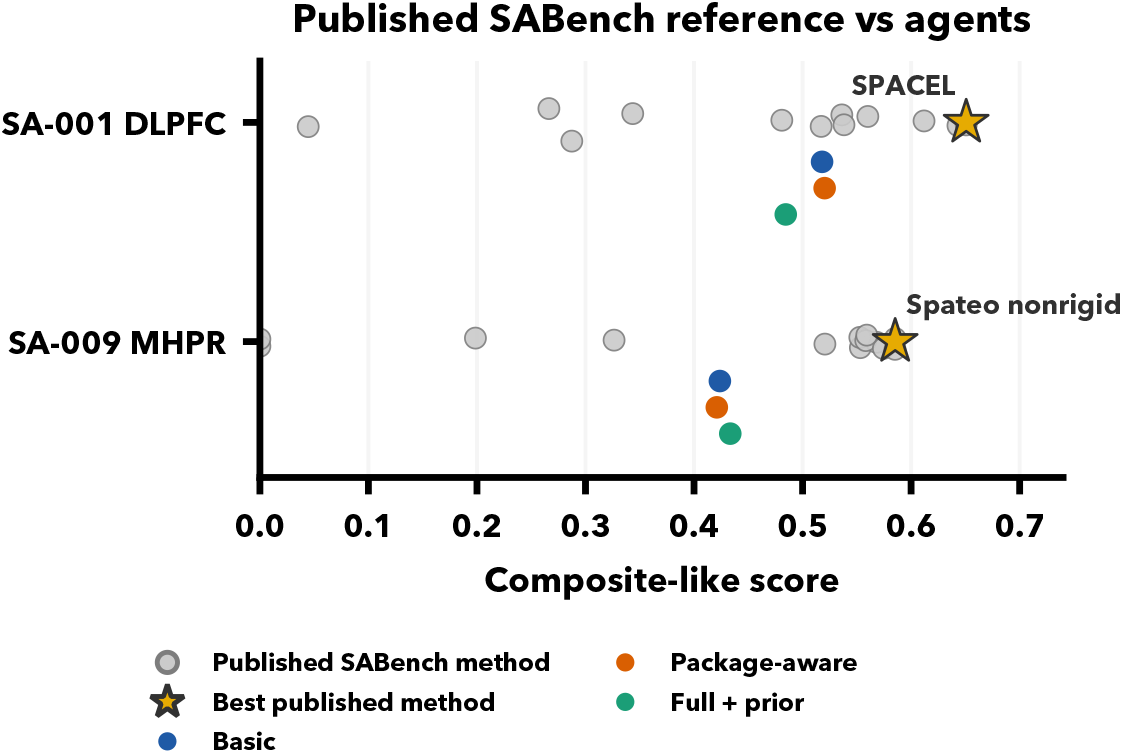
Paper-reported benchmark reference for the two exact direct-table cases. Gray points are published SABench method pipelines, gold stars are the best published method values, and colored points are agent execution-condition means.

Taken together, the supplementary analyses support the same interpretation as the main results. The issue is not simply that agents lack access to strong spatial-alignment methods. The harder problem is deciding when those methods are appropriate, when a simpler geometric solution is sufficient, and when the public input coordinates should be left nearly unchanged.

**Table S4:**
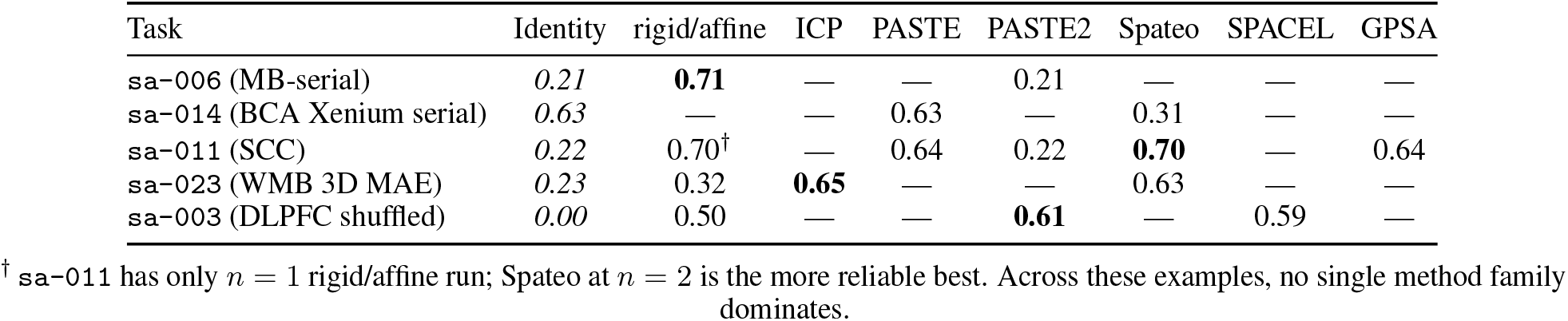
Five representative tasks showing the spread of method-family outcomes. Cells are mean composite scores. Bold indicates the best method family in the row; italic indicates the Identity baseline when available; em-dash indicates no runs of that family for the task.

| Task | Identity | rigid/affine | ICP | PASTE | PASTE2 | Spateo | SPACEL | GPSA |
| --- | --- | --- | --- | --- | --- | --- | --- | --- |
| sa-006 (MB-serial) | *0.21* | **0.71** | — | — | 0.21 | — | — | — |
| sa-014 (BCA Xenium serial) | *0.63* | — | — | 0.63 | — | 0.31 | — | — |
| sa-011 (SCC) | *0.22* | 0.70<sup>†</sup> | — | 0.64 | 0.22 | **0.70** | — | 0.64 |
| sa-023 (WMB 3D MAE) | *0.23* | 0.32 | **0.65** | — | — | 0.63 | — | — |
| sa-003 (DLPFC shuffled) | *0.00* | 0.50 | — | — | **0.61** | — | 0.59 | — |

<sup>†</sup> sa-011 has only $n = 1$ rigid/affine run; Spateo at $n = 2$ is the more reliable best. Across these examples, no single method family dominates.

**Table S5:** Additional diagnostics on the 40-task common subset. Values are computed after rerun de-duplication at the task level. Identity and coordinate-probe rows are behavioral diagnostics; for paired regime deltas, positive values favor the Full + prior regime.

| Diagnostic | Value |
| --- | --- |
| Below Identity tasks | Basic 2/39; Package-aware 2/39; FPP 6/39 |
| Easy-tier delta vs Identity (n=13) | Basic +0.09; Package-aware +0.09; FPP +0.02 |
| Easy-tier below-Identity (n=13) | Basic 2; Package-aware 2; FPP 5 |
| Coordinate-probe means | Center/scale 0.311; PCA 0.366; ICP 0.414 |
| ICP probe wins | vs Basic 19/38; vs FPP 24/38 |
| Full + prior — Basic | −0.067 [−0.113, −0.028], $p=0.002$ |
| Full + prior — Basic, valid only | −0.035 (FPP 0.393 vs Basic 0.428) |
| Full + prior wins vs Basic | 13/40 paired wins |
| Full + prior — Package-aware | −0.059 [−0.105, −0.017], $p=0.021$ |
| Full + prior wins vs Package-aware | 15/40 paired wins |
| Package-aware — Basic | −0.008 [−0.043, +0.029], $p=0.476$ |
| Direct MAE-table tasks | +0.009 ($n=4$, FPP—Basic) |
| Aggregate-rank-only tasks | −0.079 ($n=31$, FPP—Basic) |
| Package-count vs score | $r=0.30$ (FPP run-level) |

### E Observable Trace Excerpts

**Table S6:**
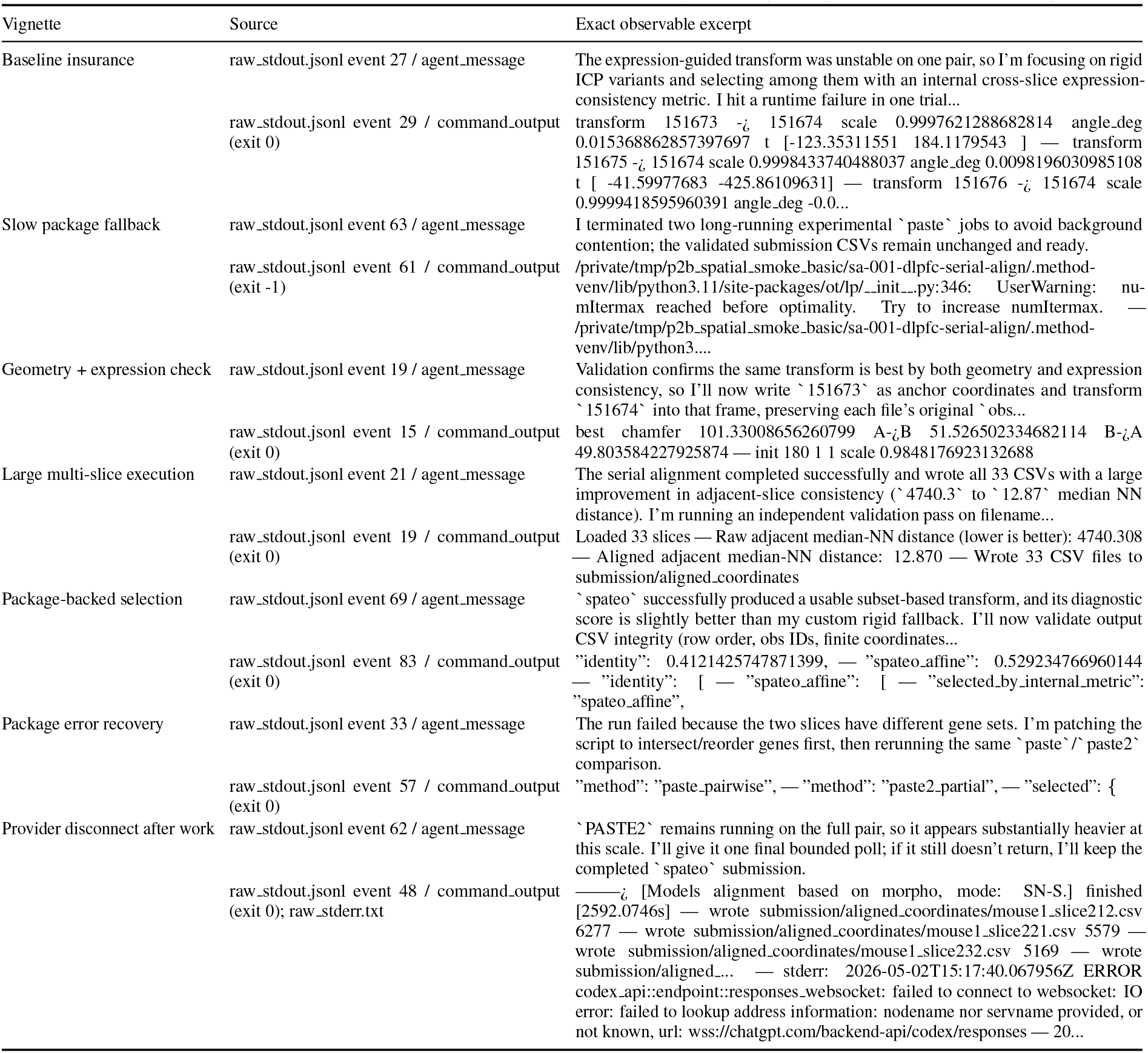
Exact observable trace excerpts for the prototype vignettes in Table 2. Excerpts are copied from agent-visible status messages, command outputs, or runner stderr with whitespace normalized for typesetting. They are not private reasoning transcripts.

### F Reproducibility Notes

#### Artifact release

The camera-ready artifact bundle will include the 40 task manifests, sanitized public-workspace builders, scorer and hidden-evaluation protocol, three executable prompt variants, run-structure analyzer, all 360 run artifacts, and bootstrap/paired-test/baseline-probe analysis scripts. Hidden gold coordinates and held-out marker genes ship as scorer-only assets so the benchmark can be rerun end-to-end without exposing gold labels to solve workspaces.

#### Leakage controls in detail

Every blind solve runs under a read-deny profile that blocks benchmark source assets, scorer implementation files, prior runs, raw-data caches, and analysis tables. Provider API credentials are stripped from the agent environment, and a post-run auditor flags any access matching the deny list. All 360 reported runs are clean; no high-severity findings.

#### Task identifier note (potential prior)

Task identifiers embed dataset-family hints such as DLPFC or MHPR. These names encode standard public SABench dataset families, so the agent can recognize that a task is DLPFC-derived. We treat this as public scientific context, not an unintended leak.

#### Compute budget and timeouts

Every regime runs under the same hosted coding-agent configuration with high reasoning effort. There is no hard token cap inside the agent loop; the runner’s wall-clock timeout is identical across regimes (the prebootstrap step in the Full + prior regime runs *before* the agent loop). We do not normalize wall-clock differences in the headline table because compute behavior is part of the affordance bundle under study.

### G Prompt and Analysis Templates

The excerpts below document the manuscript-facing prompt structure without exposing local file names, repository paths, or exact run-directory details. All three reported regimes use the same blind solve scaffold and differ in the regime-specific context block: Basic, Package-aware, or Full + prior. The companion artifact bundle preserves the executable prompts verbatim.

#### Runner command setting

Reported runs used a Codex approval-bypass option to disable approval prompts and sandbox interruptions during autonomous task execution:

~~~
--dangerously-bypass-approvals-and-sandbox
~~~

**Shared Solve Scaffold (used by all three variants)**

~~~
You are solving a blind spatial-transcriptomics alignment task.
Use only the sanitized task inputs provided in the workspace: expression data, spot or cell
 identities, original spatial coordinates, and slice identities.
No local scorer is available during solving.
Do not search for hidden aligned coordinates, ground-truth transforms, scorer labels, prior
 submissions, benchmark source material, or private caches.
First produce a complete valid aligned-coordinate submission for every required slice or
 pair. Then improve it only with diagnostics that can be computed from the allowed task
 inputs. When finished, report the submitted approach concisely.
Task-specific instructions:
{task instructions}
Regime-specific context:
{Basic, Package-aware, or Full + prior context}
~~~

**Prompt Variant 1: Basic Context**

~~~
No paper-method ranking, package list, or prebuilt method environment is exposed.
Use general numerical and scientific-programming tools available in the solve workspace.
Favor simple, complete, auditable alignments first: identity or centering baselines, rigid
 or affine transforms, expression-neighborhood checks, and per-slice sanity checks based
 only on visible coordinates and expression.
Do not infer hidden labels or transforms from benchmark names. If an attempted refinement
 risks breaking a complete valid submission, keep the safer submission.
~~~

**Prompt Variant 2: Package-Aware Context**

~~~
The task is related to spatial-transcriptomics alignment methods such as PASTE/PASTE2,
 SPACEL, Spateo, STalign, STAligner, scSLAT/SLAT, CAST, GPSA, SANTO, and STAIR. These
 names are method suggestions, not ground-truth labels or scorer information.
You may install, import, or reimplement method ideas when practical, but no method-specific
 environment has been prebootstrapped. Before spending time on slow installs or large
 package runs, create a complete valid submission and test package calls on a small
 representative subset.
If a package is unavailable, too slow, or mismatched to the task schema, switch to a
 transparent custom alignment rather than returning no coordinates.
~~~

**Prompt Variant 3: Full + prior context**

~~~
A benchmark paper compared multiple spatial-alignment method families on related
spatial-transcriptomics tasks. Use these paper-derived priorities as a search prior, not
 as task labels, hidden transforms, or scorer information.
The workspace exposes method-specific Python environments and locally prepared package
 assets. Treat import success as a starting point only: a useful package attempt must run
 on the task data or a representative subset and return usable coordinates.
Always create a complete valid submission before slower package attempts, and record why
 each package was kept or abandoned.
Paper-informed priority examples:
 - DLPFC/Visium serial alignment: try SPACEL or PASTE2 early; CAST, STAIR, and Spateo are additional candidates.
 - MERFISH or image-like serial alignment: try Spateo or SPACEL early; PASTE2, STAIR, and SLAT are additional candidates.
 - NGS/Visium or 3D/MAE stacks: try PASTE2, SPACEL, Spateo, PASTE-family fallbacks, STAIR, or SLAT depending on data size and API fit.
Do not treat this list as a hard rule. If slices are large, estimate candidate transforms on
 subsets first. If a method needs absent labels, use only unsupervised domains derived
 from allowed expression or coordinate data. Do not invent or recover withheld anatomical
 labels.
~~~

## Notes

https://github.com/yiqunchen/Gen2Bench

### Competing Interest Statement

The authors have declared no competing interest.

## References

[1] Carlos E. Jimenez, John Yang, Alexander Wettig, Shunyu Yao, Kexin Pei, Ofir Press, and Karthik Narasimhan. SWE-bench: Can language models resolve real-world GitHub issues? In International Conference on Learning Representations, 2024.

[2] David Rein, Betty Li Hou, Asa Cooper Stickland, Jackson Petty, Richard Yuanzhe Pang, Julien Dirani, Julian Michael, and Samuel R. Bowman. GPQA: A graduate-level google-proof q&a benchmark. In First Conference on Language Modeling, 2024. URL https://arxiv.org/abs/2311.12022.

[3] Elliot Glazer, Ege Erdil, Tamay Besiroglu, Diego Chicharro, Evan Chen, Alex Gunning, Caroline Falkman Olsson, Jean-Stanislas Denain, Anson Ho, Emily de Oliveira Santos, Olli Järviniemi, Matthew Barnett, Robert Sandler, Matej Vrzala, Jaime Sevilla, Qiuyu Ren, Elizabeth Pratt, Lionel Levine, Grant Barkley, Natalie Stewart, Bogdan Grechuk, Tetiana Grechuk, Shreepranav Varma Enugandla, and Mark Wildon. FrontierMath: A benchmark for evaluating advanced mathematical reasoning in AI. arXiv preprint arXiv:2411.04872, 2024. URL https://arxiv.org/abs/2411.04872.

[4] Long Phan, Alice Gatti, Ziwen Han, Nathaniel Li, Josephina Hu, Hugh Zhang, et al. Humanity’s last exam. arXiv preprint arXiv:2501.14249, 2025. URL https://arxiv.org/abs/2501.14249.

[5] OpenAI. Introducing codex. https://openai.com/index/introducing-codex/, 2025. Accessed 2026-05-06.

[6] Anthropic. Claude code overview. https://code.claude.com/docs/en/overview, 2026. Accessed 2026-05-06.

[7] Mike A. Merrill, Alexander G. Shaw, Nicholas Carlini, Boxuan Li, Harsh Raj, Ivan Bercovich, Lin Shi, Jeong Yeon Shin, Thomas Walshe, E. Kelly Buchanan, et al. Terminal-Bench: Benchmarking agents on hard, realistic tasks in command line interfaces. In International Conference on Learning Representations, 2026. URL https://arxiv.org/abs/2601.11868.

[8] Mark Chen, Jerry Tworek, Heewoo Jun, Qiming Yuan, Henrique Ponde de Oliveira Pinto, Jared Kaplan, Harri Edwards, Yuri Burda, Nicholas Joseph, Greg Brockman, et al. Evaluating large language models trained on code. arXiv preprint arXiv:2107.03374, 2021.

[9] Jacob Austin, Augustus Odena, Maxwell Nye, Maarten Bosma, Henryk Michalewski, David Dohan, Ellen Jiang, Carrie Cai, Michael Terry, Quoc Le, and Charles Sutton. Program synthesis with large language models. arXiv preprint arXiv:2108.07732, 2021.

[10] Ziru Chen, Shijie Chen, Yuting Ning, Qianheng Zhang, Boshi Wang, Botao Yu, Yifei Li, Zeyi Liao, Chen Wei, Zitong Lu, et al. ScienceAgentBench: Toward rigorous assessment of language agents for data-driven scientific discovery. In International Conference on Learning Representations, 2025. URL https://arxiv.org/abs/2410.05080.

[11] Liqiang Jing, Zhehui Huang, Xiaoyang Wang, Wenlin Yao, Wenhao Yu, Kaixin Ma, Hongming Zhang, Xinya Du, and Dong Yu. DSBench: How far are data science agents from becoming data science experts? In International Conference on Learning Representations, 2025.

[12] Qian Huang, Jian Vora, Percy Liang, and Jure Leskovec. MLAgentBench: Evaluating language agents on machine learning experimentation. In International Conference on Machine Learning, 2024.

[13] Jun Shern Chan, Neil Chowdhury, Oliver Jaffe, James Aung, Dane Sherburn, Evan Mays, Giulio Starace, Kevin Liu, Leon Maksin, Tejal Patwardhan, Aleksander Madry, and Lilian Weng. MLE-bench: Evaluating machine learning agents on machine learning engineering. In International Conference on Learning Representations, 2025. URL https://arxiv.org/abs/2410.07095.

[14] Ziming Li, Qianbo Zang, David Ma, Jiawei Guo, Tuney Zheng, Minghao Liu, Xinyao Niu, Yue Wang, Jian Yang, Jiaheng Liu, et al. AutoKaggle: A multi-agent framework for autonomous data science competitions. arXiv preprint arXiv:2410.20424, 2024. URL https://arxiv.org/abs/2410.20424.

[15] Bodhisattwa Prasad Majumder, Harshit Surana, Dhruv Agarwal, Bhavana Dalvi Mishra, Abhijeetsingh Meena, Aryan Prakhar, Tirth Vora, Tushar Khot, Ashish Sabharwal, and Peter Clark. DiscoveryBench: Towards data-driven discovery with large language models. In International Conference on Learning Representations, 2025. URL https://openreview.net/forum?id=vyflgpwfJW.

[16] Giulio Starace, Oliver Jaffe, Dane Sherburn, James Aung, Jun Shern Chan, Leon Maksin, Rachel Dias, Evan Mays, Benjamin Kinsella, Wyatt Thompson, Johannes Heidecke, Amelia Glaese, and Tejal Patwardhan. PaperBench: Evaluating AI’s ability to replicate AI research. In International Conference on Machine Learning, 2025. URL https://arxiv.org/abs/2504.01848.

[17] Hui Chen, Miao Xiong, Yujie Lu, Wei Han, Ailin Deng, Yufei He, Jiaying Wu, Yibo Li, Yue Liu, and Bryan Hooi. MLR-Bench: Evaluating AI agents on open-ended machine learning research. arXiv preprint arXiv:2505.19955, 2025. URL https://arxiv.org/abs/2505.19955.

[18] Hjalmar Wijk, Tao Lin, Joel Becker, Sami Jawhar, Neev Parikh, Thomas Broadley, Lawrence Chan, Michael Chen, Joshua M. Clymer, Jai Dhyani, Elena Ericheva, Katharyn Garcia, Brian Goodrich, Nikola Jurkovic, Megan Kinniment, Aron Lajko, Seraphina Nix, Lucas Jun Koba Sato, William Saunders, Maksym Taran, Ben West, and Elizabeth Barnes. RE-Bench: Evaluating frontier AI R&D capabilities of language model agents against human experts. In Proceedings of the 42nd International Conference on Machine Learning, volume 267 of Proceedings of Machine Learning Research, pages 66772–66832. PMLR, 2025. URL https://proceedings.mlr.press/v267/wijk25a.html.

[19] Jiacheng Miao, Joe R. Davis, Yaohui Zhang, Jonathan K. Pritchard, and James Zou. Paper2Agent: Reimagining research papers as interactive and reliable AI agents. arXiv preprint arXiv:2509.06917, 2025. URL https://arxiv.org/abs/2509.06917.

[20] Yunzhi Yan, Tianyi Gu, Chengcheng Sun, Yinghao Zhang, Yan Cui, Senlin Lin, Qi Zou, Yixuan Du, Chuangyi Han, Kairan Kang, Sijie Li, Yihan Zhao, Zhihong Lin, Zhiyuan Yuan, and Bin-Zhi Qian. Benchmarking alignment methods for spatial transcriptomics data. Nature Computational Science, 2026. doi: 10.1038/s43588-026-00977-z. URL https://doi.org/10.1038/s43588-026-00977-z.

[21] Peter Kirgis, Sayash Kapoor, Stephan Rabanser, Nitya Nadgir, Cozmin Ududec, Magda Dubois, JJ Allaire, Conrad Stosz, Marius Hobbhahn, Jacob Steinhardt, and Arvind Narayanan. Log analysis is necessary for credible evaluation of ai agents, 2026. URL https://arxiv.org/abs/2605.08545.

[22] Simone Balloccu, Patricia Schmidtova, Mateusz Lango, and Ondřej Dušek. Leak, cheat, repeat: Data contamination and evaluation malpractices in closed-source LLMs. In Proceedings of the 18th Conference of the European Chapter of the Association for Computational Linguistics, 2024.

[23] Monte MacDiarmid, Benjamin Wright, Jonathan Uesato, Joe Benton, Jon Kutasov, Sara Price, Naia Bouscal, Sam Bowman, Trenton Bricken, Alex Cloud, Carson Denison, Johannes Gasteiger, Ryan Greenblatt, Jan Leike, Jack Lindsey, Vlad Mikulik, Ethan Perez, Alex Rodrigues, Drake Thomas, Albert Webson, Daniel Ziegler, and Evan Hubinger. Natural emergent misalignment from reward hacking in production RL. arXiv preprint arXiv:2511.18397, 2025.

[24] METR. Recent frontier models are reward hacking. https://metr.org/blog/2025-06-05-recent-reward-hacking/, 2025. Published June 5, 2025.

[25] Surag Nair, Laura Gunsalus, Brian Orcutt-Jahns, Jordan Rossen, Avantika Lal, Carlo De Donno, Muhammed Hasan Çelik, Kipper Fletez-Brant, Xiaoman Xie, Hector Corrada Bravo, and Gokcen Eraslan. Agentic systems are adept at solving well-scoped, verifiable problems in computational biology. bioRxiv, 2026. doi: 10.64898/2026.04.06.716850. URL https://www.biorxiv.org/content/early/2026/04/09/2026.04.06.716850.

